# Intramolecular dynamics of single molecules in free diffusion

**DOI:** 10.1101/120311

**Authors:** Charles Limouse, Jason C. Bell, Colin J. Fuller, Aaron F. Straight, Hideo Mabuchi

## Abstract

Biomolecular systems such as multiprotein complexes or biopolymers can span several tens to several hundreds of nanometers, but the dynamics of such “mesocale” molecules remain challenging to probe. We have developed a single-molecule technique that uses Tracking Fluorescence Correlation Spectroscopy (tFCS) to measure the conformation and dynamics of molecular assemblies specifically at the mesoscale level (~100-1000 nm). tFCS is non-perturbative, as molecules, which are tracked in real-time, are untethered and freely diffusing. To achieve sub-diffraction spatial resolution, we use a feedback scheme which allows us to maintain the molecule at an optimal position within the laser intensity gradient. We find that tFCS is sufficiently sensitive to measure the distance fluctuations between two sites within a DNA molecule separated by distances as short as 1000 bp. We demonstrate that tFCS detects changes in the compaction of reconstituted chromatin, and can assay transient protein mediated interactions between distant sites in an individual DNA molecule. Our measurements highlight the impact that tFCS can have in the study of a wide variety of biochemical processes involving mesoscale conformational dynamics.

## Introduction

The regulation of gene expression, intracellular transport, replication, and many intracellular processes are mediated by the dynamics of molecular systems occurring at the mesoscale level (~100 – 1000 nm). Yet, methods for probing dynamics of mesoscale molecules are sparse. Super-resolution microscopy techniques [1] and biochemical methods to identify proximal interactions *in vivo* such as HiC [2, 3] BioID [4] or APEX [5], are powerful tools which can provide an equilibrium view of mesoscale organization inside the cell. However, it would be highly valuable to complement these methods with approaches that can i) report on dynamic changes in the conformation of mesoscale systems, and ii) be applied to reconstituted systems, where the dynamics of individual molecules can be measured in isolation or with selected partners, and biochemical and biophysical models can be directly tested. Single molecule methods that have been used to study mesoscale systems include tethered particle motion [6, 7] and force spectroscopy [8, 9], but these tools require physical immobilization of the molecules which may perturb their dynamics. We have developed a new approach that overcomes some of these existing limitations and enables the measurement of conformational fluctuations in mesoscale biological molecules in tether-free and force-free conditions.

Tracking-fluorescence correlation spectroscopy (tFCS) is a method which combines confocal microscopy, feedback-based single-molecule tracking, and fluorescence correlation spectroscopy (FCS) to measure the conformational dynamics of individual molecules without the need for mechanical tethering [10, 11, 12, 13, 14]. The key element in tFCS is the use of active feedback to compensate for center of mass diffusion by repositioning the microscope stage in real-time so that the fluorescence from an individual molecule can be monitored over a long time period. However, tFCS had been restricted to very large molecules such as λ-phage DNA [15] (radius of gyration ~1 μm) due to spatial resolution limitations. In parallel, other active tracking methods [16, 17, 18] have been used by several researchers to measure hydrodynamic mobilities [19, 20], stoichiometry of molecular complexes [21], or nanoscale conformational dynamics [22, 23], but none have permitted mesoscale intramolecular dynamics measurements.

Here, we developed a dual-color tFCS assay which specifically assays the distance fluctuations between two discrete sites within a single macromolecule with approximately 100 – 150 nm spatial resolution and sub-millisecond temporal resolution. By optimizing the focal position of the lasers used for tracking and intramolecular dynamics detection, we showed that tFCS has sufficient spatial and temporal resolution to probe the relative motion between two sites in freely diffusing DNA molecules separated by as little as 1000 bp. Additionally, we showed that tFCS can detect conformational transitions in diffusing chromatin fibers and DNA looping processes induced by Lac repressor. These measurements provide the first demonstration that tFCS can measure dynamic biophysical processes occurring on a sub-diffraction scale, which has important implications for many molecular systems, in particular the dynamics of nucleoprotein systems.

## Results

### Design of the tFCS assay to measure the intramolecular dynamics of freely diffusing single molecules

To measure the conformation of biological macromolecules in absence of mechanical perturbation, we custom-designed a confocal microscope with a feedback system that compensates for the motion of individual molecules as they freely diffuse [13]. This makes it possible to track single molecules in three-dimensional space while simultaneously observing their fluorescence emissions (Fig. 1A). We tailored the system to specifically assay the distance and distance fluctuations between two parts of a single molecule or complex. To accomplish this, we used a green reference dye (Cy3b) as a reference point for tracking the movement of the molecule in three dimensions, and a red probe dye (Atto647N) as a reporter of the intramolecular dynamics. We then controlled, via feedback, the position of two lasers in real-time to simultaneously excite the reference tracking and probe dyes. To follow the diffusive path of the reference dye, we rapidly modulated the 3D position of the tracking laser to generate a position sensitive fluorescence signal [11, 13, 24]. Importantly, we coordinated the positions of the probe and tracking lasers (Fig. 1B) to effectively maintain the probe laser focus at a fixed point relative to the reference site of the molecule. Using this labeling strategy which employs only two fluorophores, we were able to track individual molecules for several seconds, and collect >1 million photons per molecule in both fluorescence channels (**Fig. S1, Table S2**).

**Figure 1.**
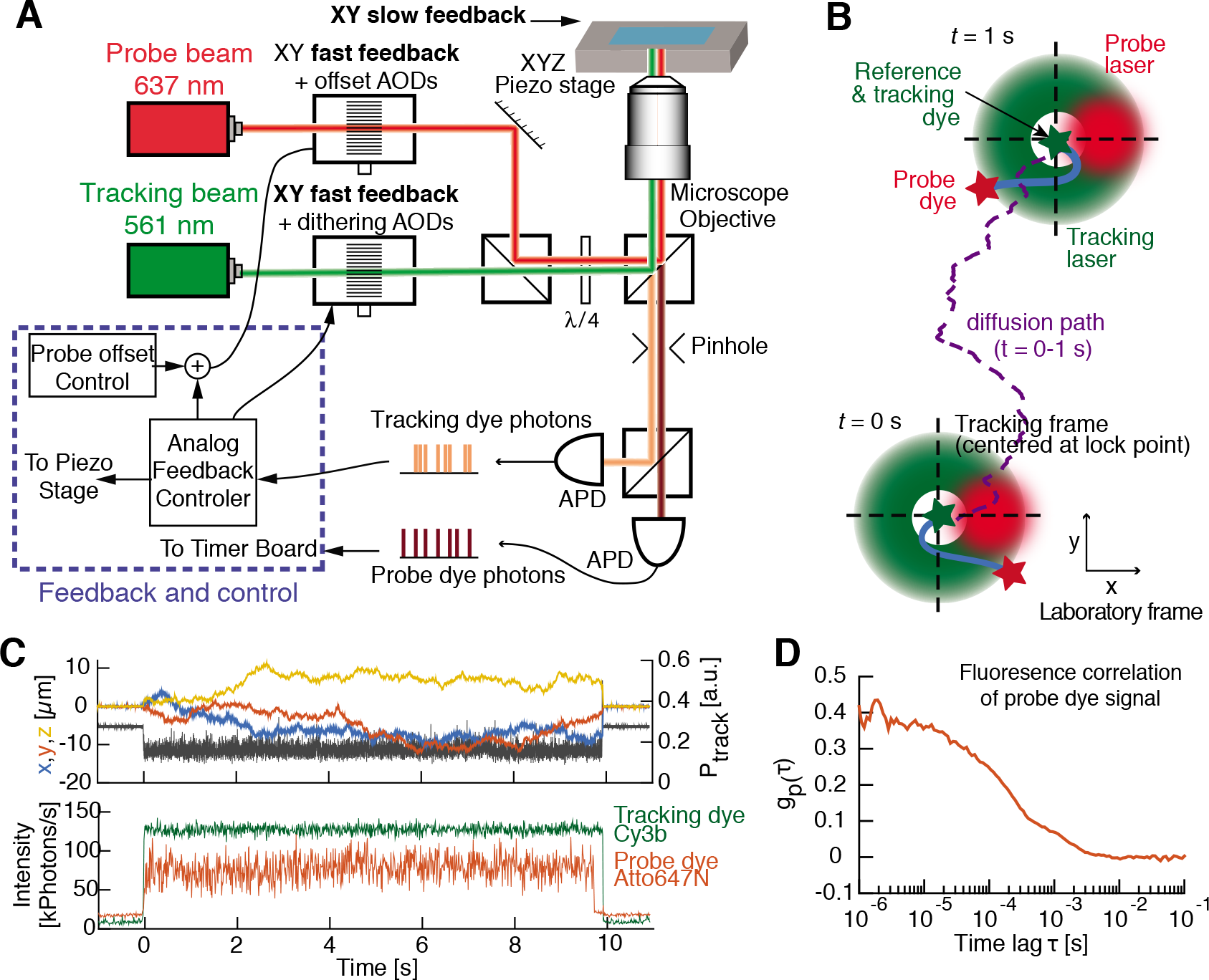
Instrumentation and concept of the tFCS assay. (**A**) Key optical and electronic elements of the tFCS microscope. The feedback is applied by an analog controller (blue box), which drives the microscope piezo stage and the acousto-optic deflectors (AODs) controlling the position of the tracking and probe beams. (**B**) Schematic of the tracking and probe beam motions during tracking of a freely diffusing molecule (blue). The trajectory (purple line) of the green labeled site on the molecule (green star, reference dye) is followed by the tracking laser (green beam), while the probe laser (red beam) is maintained at a fixed position with respect to the tracking laser. The green labeled site is immobile with respect to the probe laser (at the tracking/reference lock-point, dashed crosshair), while the red labeled site (red star, probe dye) moves due to intramolecular dynamics. Green ring indicates circular dithering of the tracking beam used to locate the molecule [13]. (**C**) Single-molecule signals recorded in tFCS, including the probe fluorescence (red), the stage trajectory, and the tracking laser power (black) which is under feedback control to keep the tracking dye fluorescence at a fixed value (green). (**D**) Single-molecule fluorescence correlation function *g_p_*(*τ*) from the probe signal shown in (C).

The schematic representation of a typical trajectory of the molecule and the focal positions of the lasers (Fig. 1B) highlights how intramolecular dynamics gives rise to fluctuations of the probe dye fluorescence due to changes in either the distance or orientation between the probe and reference sites. To quantify the reference-to-probe dye dynamics, we computed the autocorrelation of the probe dye fluorescence signal as is standard in conventional FCS (Fig. 1C, D). From the three-dimensional movement of the molecule over time provided by the *X, Y, Z* stage trajectory (Fig. 1C), we obtained an independent and calibrated readout of the diffusion coefficient. Our approach is generalizable to any biological macromolecule that can be labeled with two different fluorophores. The labeling sites can be located on the same individual molecule or on two separate binding partners. Thus, by selecting the appropriate position of labeling sites, we can measure any desired pairwise distance within a molecule or macromolecular complex.

### Optimization of the spatial sensitivity of tFCS through optimization of the illumination geometry

A major challenge for mesoscale measurements is that the target resolution (~100 – 1000 nm) is smaller than the size of the diffraction limited probe beam. In prior implementations of tFCS, the intramolecular motion was larger than the beam dimensions, and the fluorescence fluctuations stemmed from large excursions of the probe dye in and out of the entire Gaussian beam profile [15]. In this new tFCS assay, the trajectory of the probe dye samples only a small region of the illumination volume. To resolve intramolecular dynamics in the sub-diffraction-limit regime, we increased the spatial sensitivity of the assay by configuring the feedback to maintain the molecule at the edge of the Gaussian profile of the probe laser, where the local intensity gradient is large (Fig. 2A, B). If the reference and probe beams were collinear (centered-illumination), the local intensity gradient seen by the probe fluorophore would be small and intramolecular dynamics would give rise to small fluorescence fluctuations (Fig. 2A). Therefore, we modified the microscope optical path so that the probe beam was laterally offset with respect to the tracking beam. We tuned the distance between the probe and tracking beam electronically using the acousto-optic Deflectors (AODs) in the probe beam path (Fig. 1A). With this “side-illuminatio” configuration, even small displacements of the probe dye result in large changes in the laser intensity seen by the probe dye, which provides a sensitive way to convert molecular motion into fluorescence fluctuations (Fig. 2B, **S2**).

**Figure 2.**
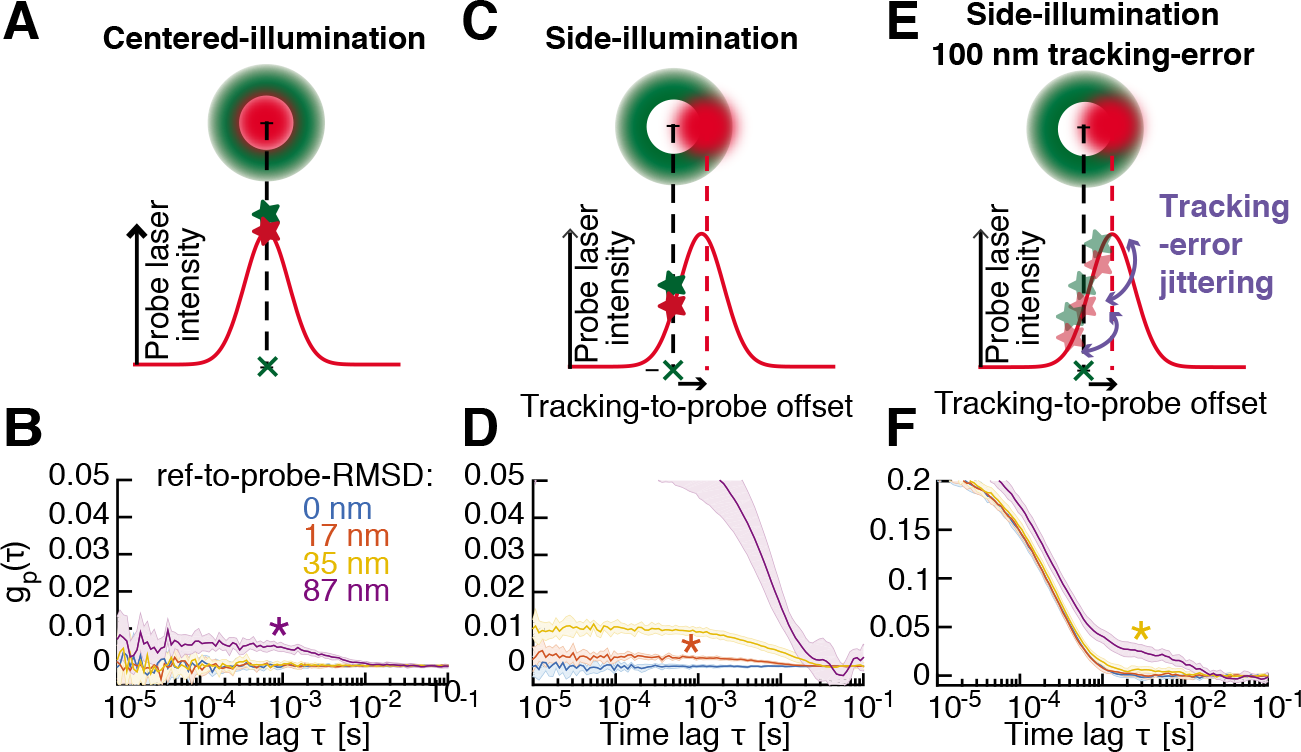
Optimization of tFCS spatial sensitivity with side-illumination, and expected resolution in presence of tracking localization error. (**A, C**) Schematic of the centered- (A) and side-illumination (C) configurations. Top: Relative positions of the tracking (green) and probe (red) beams. Bottom: *X*-axis cross-section of probe laser intensity profile seen by the probe dye. Green cross represents the lock-point of the feedback (at the center of tracking laser circular orbit) with respect to which the reference dye is immobile. Green and red stars represent the average position of the reference and probe dyes in the beam, respectively. (**B, D**) Simulated tFCS signals (mean ± 95% CI) of molecules with a reference-to-probe dye root mean squared distance of 0, 17, 35 or 87, using either the centered- (B), or the side-illumination (D) configuration. In the side-illumination geometry, the reference-to-probe beam offset was set to 1.0w, where *w* = 310 nm. Asterisk color indicates the shortest resolvable reference-to-probe RMSD. (**E**) Illustration showing the effect of finite-bandwidth tracking, where the reference dye is imperfectly maintained at the lock-point, which results in a residual jitter of the probe dye in the probe beam frame. (**F**) Same simulation as in (D), but taking into account imperfect feedback (100 nm RMS localization error).

### Effect of tracking-localization error and experimental considerations for fast feedback

To obtain a high spatial resolution, it is essential that the microscope feedback loop tracks the displacement of the reference dye as accurately as possible. In practice, the actuators bandwidth and the finite dye brightness set limits on the feedback bandwidth and tracking accuracy [11]. This leads to a residual motion of the probe dye in the probe beam which impairs spatial sensitivity as it is equivalent to random displacements of the probe beam away from its optimal position (Fig. 2C) and reduces the local intensity gradient seen by the probe dye (Fig. 2G).

To maximize tracking accuracy, we improved the feedback architecture so that we could track the molecule with a bandwidth larger than the resonance frequency of the piezo stage (~ 100 Hz). To do so, we implemented, in the *XY* dimensions, a feedback loop with two branches. We corrected for low-frequency components of the particle motion via feedback on the microscope piezo stage, whereas higher bandwidth components were canceled via feedback on the laser position, controlled with AODs. With this scheme we were able to feedback at ~1 kHz bandwidth for *XY* tracking and maintain the reference dye within better than 100 nm of the desired lock point (RMS error) for molecules diffusing up to 15 μm^2^/s.

### Numerical simulations and estimation of the spatial resolution of tFCS

To test whether our tFCS method can resolve intramolecular dynamics at the mesoscale level, we first conducted numerical simulations. We simulated tFCS signals for molecules with a reference-to-probe dye root mean square distance (RMSD) ranging from 17 to 87 nm (Fig. 2D, E, F), and a characteristic timescale of intramolecular motion of 10 ms. We chose simulation parameters (number of photons collected, signal-to-noise, duration of traces) which matched typical experimental values (**Fig. S1, Table S2**). We then compared the tFCS signal obtained for each reference-to-probe RMSD with that of control molecules where the reference and probe sites were co-localized (i.e., reference-to-probe RMS distance equal 0). In the ideal case of perfect tracking, the side-illumination configuration would allow us to resolve the intramolecular dynamics of all the molecules ranging from 17 nm and upward (Fig. 2D), compared with the centered-illumination geometry, where only those molecules with a reference-to-probe RMS distance of 87 nm would be resolved (Fig. 2E). When we accounted for a realistic tracking localization RMS error of 100 nm, this reduced the resolution to 35 nm with the side-illumination configuration (Fig. 2H,I). Simulations with different intramolecular timescales of motion (1 ms and 10 ms) and different offset values in the side-illumination configuration yielded similar results (**Fig. S3**). Together, these results suggested that our tFCS technique with the current performance of our tracking-microscope is sensitive to mesoscale dynamics. Additionally, the spatial resolution could be pushed well below 100 nm with a smaller tracking localization error, which can be done by using brighter fluorophores [25, 26].

### Measurement of the intramolecular dynamics of DNA

To test the ability of our approach to resolve the conformational fluctuations of macromolecules, we applied tFCS to measure the end-to-end dynamics of freely diffusing DNA molecules. Previous single-molecule studies of DNA Brownian dynamics have used ~50 kbp long polymers of λ-phage DNA [15, 27]. In our assay, we measured the end-to-end dynamics of DNA molecules one to two orders of magnitude shorter than those previously studied. We labeled dsDNA of about 0.5 kbp, 1 kbp and 3.9 kbp with a single Cy3b and a single Atto647N at opposite ends (OE) of the molecule. As a control, DNA of identical lengths were labeled with the two dyes placed on the same end of the duplex (SE), separated by only 31 bp.

We clearly resolved the end-to-end dynamics of both the 1 kbp and 3.9 kbp DNA when we used the side-illumination geometry (Fig. 3A). Conversely, we found that the centered-illumination geometry with near-perfect alignment of the probe and tracking beams resulted in poorer sensitivity, and could only resolve the 3.9 kbp DNA (Fig. 3B). The 0.5 kbp DNA fragment was below the resolution threshold for both configurations, as suggested by the overlap of the correlation signal of the SE and OE labeled molecules. When observed with the side-illumination geometry, the 3.9 kbp fragments exhibited fluorescence fluctuations with larger correlation amplitude at all time-lags and slower timescales of motion compared with the 1 kbp molecules, consistent with a larger radius of gyration and slower polymer relaxation modes. Even though the reference and probe sites were nearly co-localized in the same-end labeled DNA molecules, they exhibited a clearly non-zero fluorescence correlation signal (Fig. 3A, B) indicating the presence of systematic noise in the tFCS assay. This systematic noise likely stems from imperfect localization of the reference dye as inferred from our previous simulations (Fig. 2H, I), and from the triplet-state dynamics of the Atto647N dye, also observed in conventional FCS measurements [28, 29].

**Figure 3.**
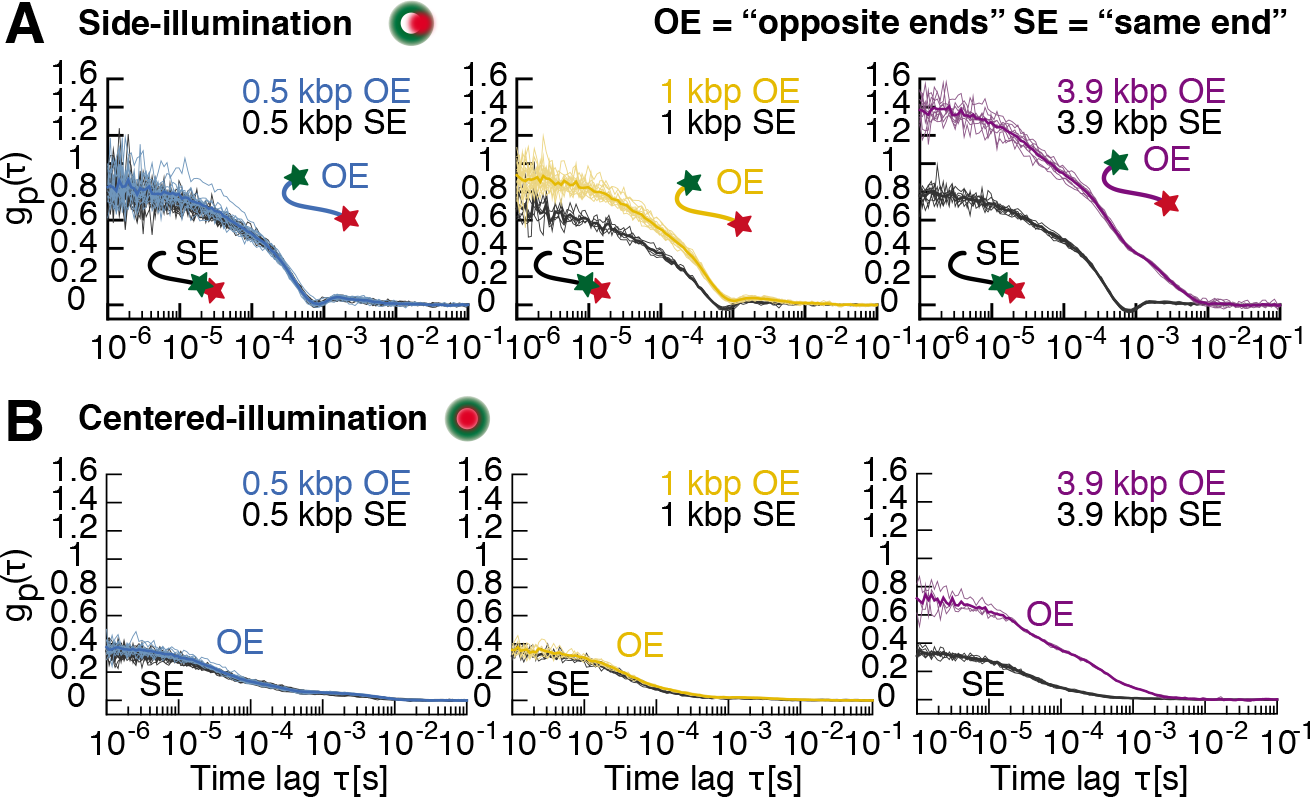
tFCS measurement of the end-to-end dynamics of freely diffusing DNA molecules. (**A, B**) Fluorescence correlation signals of DNA molecules using the side- (A) or centered-illumination (B) configuration. DNA were labeled with Cy3b and Atto647N either on opposite ends (OE) or on the same end (SE) of the molecule. DNA lengths were 0.5 kbp (blue OE, black SE), 1 kbp (yellow OE, black SE) and 3.9 kbp (purple OE, black SE). Correlation signals from individual molecules (thin lines) and population averages (thick lines) are shown.

To test the effect of changing the relative position of the reference and probe lasers, which results in a change in the local laser intensity gradient seen by the probe dye, we repeated the measurement with varying tracking-to-probe beam offsets. We found that the changes in the amplitude of the correlation signal and of the mean fluorescence signal had a quadratic dependency in the offset, as predicted analytically (**Fig. S4, SI Appendix**). Together, these results demonstrate that the tFCS assay detects the end-to-end dynamics of DNA constructs of ~1 kbp and longer, consistent with an ability to resolve intramolecular motion on the order 100-150 nm [30]. To our knowledge, this is the first single-molecule measurement of freely diffusing DNA dynamics in the semi-flexible polymer regime. Importantly, the assay separates molecules that differ only in the distance between the reference and probe sites (SE and OE constructs) and not in their overall dimensions, indicating that this is an appropriate tool for measuring macromolecular conformation in addition to its ability to measure hydrodynamic properties.

### Measuring chromatin compaction in freely diffusing molecules

Changes in the local compaction of chromatin regulate multiple essential processes including gene expression, mitotic chromosome condensation and heterochromatin formation [31, 32, 33]. Reconstituted arrays of nucleosomes [34] can recapitulate the mechanics of chromatin condensation in response to electrostatic forces [35, 36], chromatin binding proteins [37], and histone modifications or variants [38]. Traditionally, measurements of chromatin compaction in solution use sedimentation assays or analytical ultracentrifugation (AUC) to differentially fractionate compact chromatin [35]. However, these assays require a large amount of material, do not reveal the dynamics of the folding process, and are not typically compatible with complex biochemical systems such as cellular extracts. We investigated the applicability of tFCS in characterizing the conformational state of single reconstituted nucleosome arrays and as an alternative approach to studying chromatin dynamics.

To measure the dynamics of chromatin fibers, we reconstituted nucleosomes on a tandem DNA array of 19 copies of a high affinity nucleosome positioning sequence (19x601) [39] that we labeled on either end with Cy3b and Alexa647. The nucleosome arrays exhibited a small, but significant down shift in the side-illumination tFCS signal compared with that of bare DNA, indicating a smaller distance between the two ends of the molecule as a result of DNA wrapping by the histones. Upon addition of 1.5 mM Mg^2+^, we observed significant compaction of the nucleosome arrays, which is consistent with previous studies on the folding behavior of nucleosome arrays in the presence of divalent cations [35, 40].

To further quantify the folding state of individual arrays, we computed for each molecule a compaction score quantifying the reduction in amplitude of its correlation signal with respect to the bare 19x601 DNA. We found that the compaction score was positively correlated with the diffusion coefficient (Spearman’s *χ* = 0.76, p-value *<* 10^−6^) (Fig. 4B, C), confirming that both quantities reflect on the folding state of the molecule. Interestingly, the positive correlation held true when we considered only the nucleosome arrays in the absence of Mg^2+^ (Spearman’s *χ* = 0.30, p-value *<* 0.01) or only the arrays in the presence of Mg^2+^ (Spearman’s *χ* = 0.58, p-value *<* 10^−5^), but not when we considered only bare DNA molecules (p-value = 0.74). This result suggests that the tFCS assay resolves compaction differences within the population of arrays, and that the heterogeneity in compaction scores reflects true sample heterogeneity, likely stemming from differences in the number of octamers assembled on each molecule, rather than measurement noise. Together, these observations show that the tFCS approach, in contrast with the ensemble measurement of chromatin compaction by sedimentation, allows us to measure solution changes in chromatin compaction and dynamics at the single molecule level.

**Figure 4.**
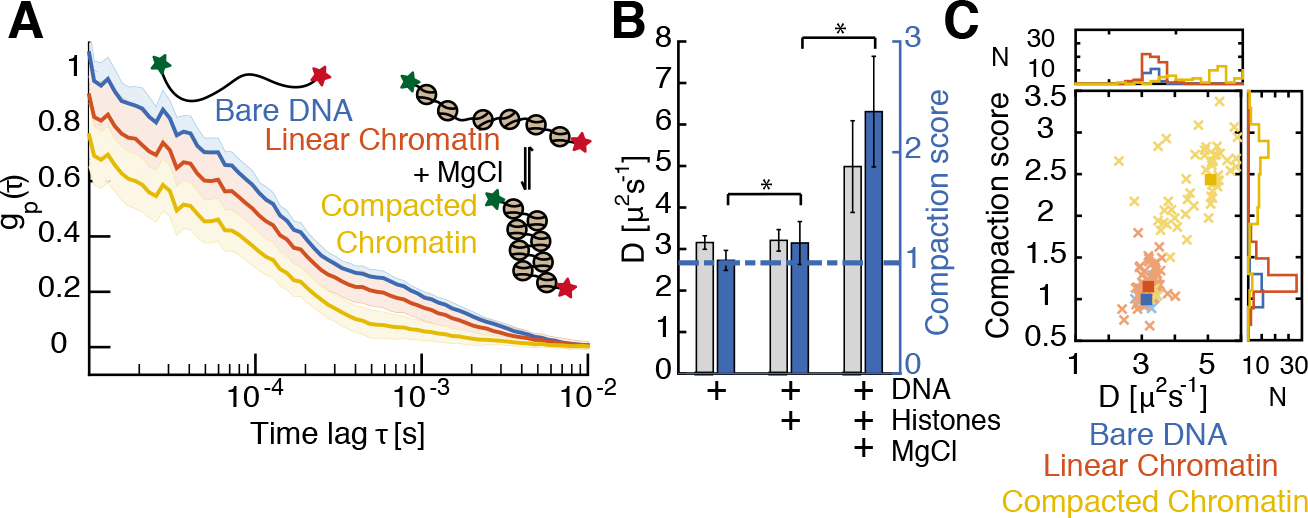
Measurement of nucleosome arrays compaction state. (**A**) tFCS signals (population mean ± std) for 19×601 DNA molecules prior to nucleosome assembly (blue), and for chromatinized 19×601 molecules in the absence (red) or presence (yellow) of 1.5 mM Mg^2+^. Data were taken using a side-illumination configuration. (**B**) Bar plot showing the mean and standard deviation of the diffusion coefficient (gray bars) and the compaction scores (blue bars) for all three samples. Asterisk indicates statistical significance (Two-sided Wilcoxon rank sum test, p-value <0.05). (**C**) Scatter plot of the diffusion coefficients and compaction scores for the three samples (same color code as in (A)). Individual molecules (light crosses) and population median (bright squares) are shown, as well as marginal distributions (top and right plots).

### Measurement of repressor induced DNA looping dynamics

Control of bacterial operons [41, 42], promoter-enhancer interactions in eukaryotes [43] and the large scale organization of topological domains in metazoan chromosomes [44] are all regulated by chromatin folding to juxtapose DNA sites separated by tens to millions of DNA bases. To demonstrate the ability of tFCS to monitor transient protein mediated interactions between remote DNA loci, we turned to the Lac operator-repressor system and asked whether we could detect the formation of looped states in freely diffusing DNA induced by Lac repressor binding [6, 41, 45] (Fig. 5).

**Figure 5.**
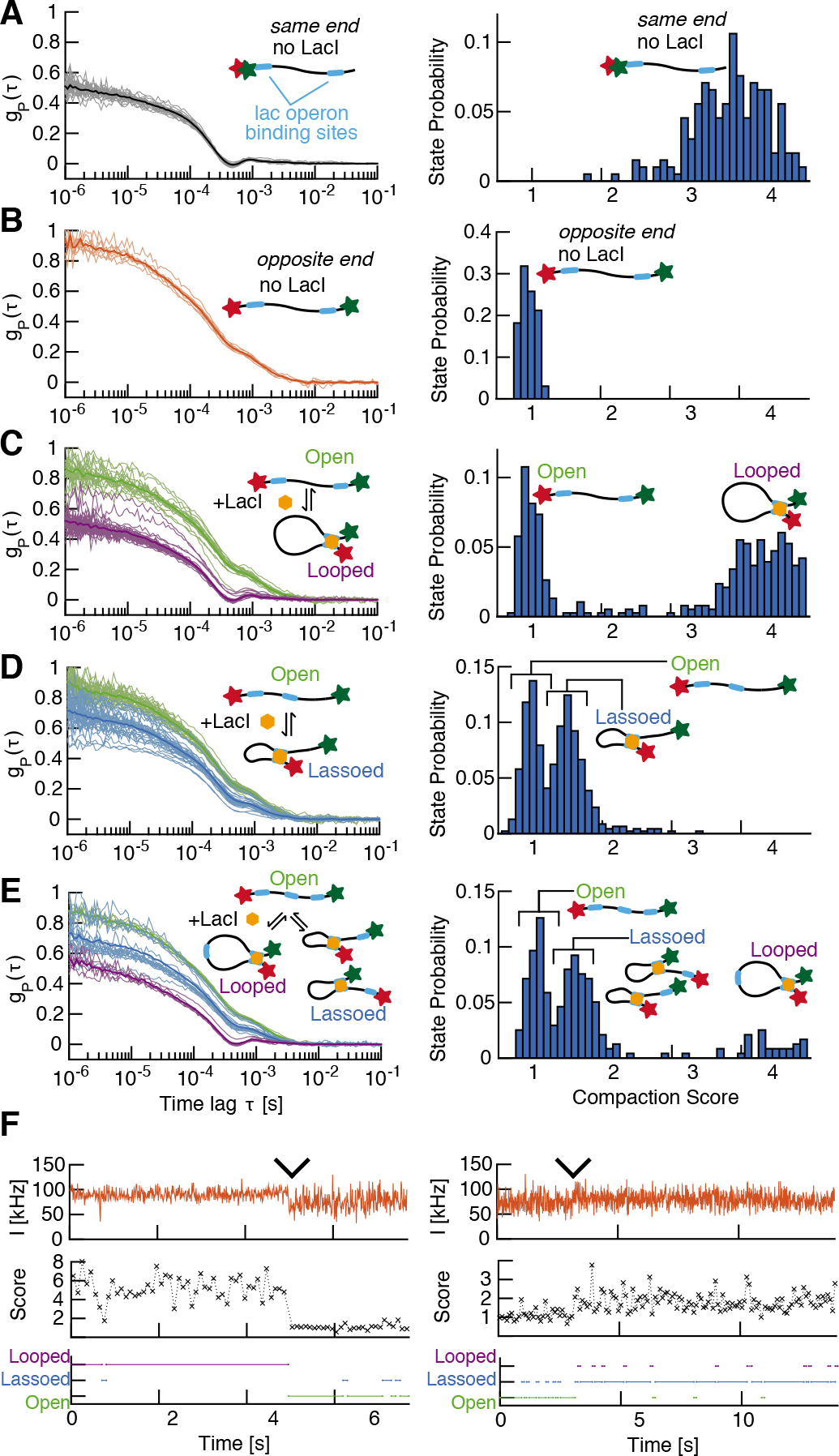
Detection of protein mediated conformational changes. (**A, B**) Left: Single-molecule tFCS signals (thin lines) and population mean (thick lines) of the looping DNA construct bearing two lac operators and labeled with Atto647N and Cy3b at the same end (SE) in (A), and at opposite ends (OE) in (B), in the absence of LacI. Right: Distribution of compaction scores for the 100 ms sub-traces of all the individual molecules shown on the left. (**C, D, E**) Left: tFCS signals of individual DNA molecules after addition of 64 nM LacI. DNA used were the looping construct in (C), the lasso construct in (D), and the construct with three LacO sites in (E). The conformational state inferred by clustering the tFCS signals are categorized as: open in green (no LacI bound or LacI bound at a single site), lassoed in blue (LacI mediated short loop), or looped in purple (LacI mediated long loop). Cluster averages (thick lines) and individual molecules (thin lines) are shown for each of these three groups. Right: Distribution of compaction scores for the 100 ms sub-traces of all the individual molecules shown on the left. (**F**) Representative example of a conformational transition between a looped and an open state (left), or a lassoed and an open state (right), detected during the tracking of an individual molecule. Probe dye fluorescence trace (top), 100 ms binned compaction score (middle), and inferred conformational state (bottom). Wedges indicate time of transition between the two conformations.

We generated two 2.6 kbp DNA substrates containing two lac operator sites spaced at specific distances (Table S1): a “looping” construct with the two binding sites at each end of the molecule, and a “lasso” construct with one operator site moved to the middle of the molecule (1.3 kbp between the operator sites). For both constructs, we found that addition of Lac repressor resulted in the appearance of two populations of molecules characterized by distinct correlation functions (Fig. 5C, D). The tFCS signal of the population with the largest correlation amplitude matched the signal measured prior to the addition of LacI (Fig. 5B), which is consistent with molecules in an unlooped conformation (with or without LacI bound). In addition, for the “looping” construct, the tFCS signal of the molecules with the smallest amplitude overlapped with the signal obtained from control DNA molecules where the two dyes were juxtaposed, validating that these molecules were in a state with the Cy3b and Atto647N dyes in close proximity (Fig. 5A). For the lasso construct, we found that the folded molecules exhibited a signal of intermediate amplitude, consistent with the formation of a short loop where the dyes are separated by about half the length of the molecule. Importantly, the addition of 2 mM of the LacI inhibitor IPTG efficiently destabilized the loops (**Fig. S5**). To test whether we could distinguish more than two conformational states, we then generated a construct containing three Lac Operator sites, one at each end of the DNA and one in the intermediate lasso position. We clearly detected the formation of both large and small loops in this construct (Fig. 5E). Altogether, these measurements demonstrate that the tFCS signal provides a single-molecule readout of the long-range interactions between specific sites, induced by LacI.

Finally, we asked whether we could detect transitions between looped and unlooped states in realtime. The LacI mediated loop lifetime (tens of seconds) is long compared with the duration of the tracking traces [6, 45]. However, using the tFCS signal computed across a 100 ms sliding window, we captured transitions in a fraction of the observed individual molecules. We compared the sliding tFCS signal with the average correlation signal of the molecules in the open, folded loop and folded lasso state, so as to infer the state of the molecule at each time point. Representative examples of an unlooping event in the loop construct (resp. looping event in the lasso construct) showed that the transitions points uncovered by classifying the 100 ms tFCS signals aligned as expected with abrupt changes in the compaction score and with visible transitions in the amplitude of fluorescence fluctuations (Fig. 5, **S6**).

## Discussion

A major challenge for studying subcellular biological processes is that few methods can interrogate the structure and dynamics of large macromolecules and complexes at the distance and time scales relevant to their function. We show here that our approach enables measurement of the internal dynamics of freely diffusing mesoscale molecules in three different experimental systems: DNA molecules in solution, chromatin fibers, and DNA looping by repressor proteins. Our method extends existing techniques such as sedimentation to the single molecule level and provides the ability to measure structural rearrangements and dynamic fluctuations that are not accessible through electron microscopic or atomic force microscopic approaches. Because our approach can be applied to freely diffusing molecules, it circumvents the pitfalls of surface tethering, which can alter the dynamic behavior of molecules. The assay is, in its aim, similar to single-molecule FRET, but while FRET is powerful for studying fast structural dynamics at the nanometer scale, it cannot be applied to distances greater than ~10 nm [46] due to intrinsic limitations in the energy transfer mechanism.

We have achieved a spatial resolution of ~10-100 nm in our current tFCS setup, however this is not a fundamental limit for the technique, but results from practical limitations in the implementation. Even with a 100 nm tracking localization RMS error, our simulations suggest a spatial resolution of ~35 nm. Anisotropic tracking accuracy, where the axial particle localization is poorer than the lateral localization resulting in suboptimal positioning of the molecule within the probe beam likely accounts for much of the discrepancy between the numerical and experimental resolution. This could be overcome with alternative optical designs, for example using a pair of perpendicularly positioned objectives, which would permit the use of acousto-optics to displace the beam along three directions of space and increase the feedback bandwidth. Imperfect coordination in the motion of the tracking and probe beams may also explain some of the discrepancy, but could be resolved with improved feedback schemes.

Fundamentally, tFCS spatial and temporal resolution are limited by photon counting in two ways: i) the fluorescence intensity of the probe dye determines the level of shot noise in the correlation signal, and ii) the count rate of the tracking dye fluorescence defines the maximum feedback bandwidth which sets the tracking accuracy. tFCS would therefore doubly benefit from brighter and more photostable tags, which are actively being developed [25, 26]. In addition, such improvements would likely enable the use of tFCS in more complex biochemical environments such as in cellular extracts, or *in vivo*.

In contrast with traditional localization microscopy techniques, spatial resolution in tFCS is not directly affected by the dimensions of the point spread function (PSF), but rather by the steepness of the laser intensity gradient at the tracking lock-point. This specificity defines an optimization problem for achieving high resolution imaging which is distinct from conventional PSF engineering. It would be interesting to investigate tFCS signals under different excitation profiles such as those used in light-sheet microscopy [47] or Stimulated-Emission-Depletion (STED) microscopy [48].

While we showed here that the intramolecular dynamics contribute to the fluorescence correlation signal, the tFCS signal also depends on the triplet-state dynamics of the probe dye, and on the residual motion of the reference dye due to tracking localization error, which makes direct fitting of the fluorescence correlation function (as done in conventional FCS) challenging. Future work will focus on deriving methods to better quantify tFCS data, with the ultimate goal of converting the fluorescence correlation signal into a distance correlation signal in physical units and infer, for example, the distance between two sites expressed in nm.

We believe that tFCS has the potential to play an important role in the biophysical realm of large molecules or complexes, similar to the role played by single molecule FRET in the study of small individual molecules (<10 nm, [49]). tFCS is applicable to probe conformational changes within any large molecular system, such as the macromolecular machines involved in transcription, replication or chromosome segregation, or within the large elongated proteins implicated in membrane transport. Our experiments highlight the usefulness of tFCS in monitoring long range contacts in DNA or more generally to characterize biophysical properties of DNA or chromatin, and may therefore be impactful in the study of genetic regulation mechanisms involving chromatin folding. We envision tFCS will be powerfully combined with biochemical manipulations, and possibly applied inside living cells, to tease apart the basic mechanisms that drive organization at the mesoscale.

## Methods

### Single molecule tracking microscope

To track individual molecules, we used a custom-built confocal microscope augmented with a feedback loop as well as beam steering capabilities that enabled high bandwidth electronic control of the position of the focus of each laser beam in the sample (details in **SI Materials and Methods**). Briefly, to achieve all-optical sensing of the 3D position of the diffusing particle and real time tracking, we set up the tracking laser according to the optical and feedback scheme previously described ([13]). To increase the tracking bandwidth of the instrument, we used two pairs of acousto-optic deflectors (AODs) placed in the path of the tracking and probe beams (Fig. 1A), to serve as fast actuators, with μs scale response time, of the transverse position of the beams. This strategy allowed us to apply feedback faster than the mechanical resonance frequency of the piezo stage (~100 Hz) and minimize the tracking localization error, which is key to resolve intramolecular dynamics at the sub-diffraction limit scale.

### Measurements conditions

All tFCS measurements were taken with molecules at around 1 pM concentration, so that so that one individual molecule drifted in the microscope confocal volume every ~15-30 s, making it unlikely to track two molecules at once. Buffer used in all experiments were 10 mM Tris pH 7.4, 50 mM KCl, ± 2 mM MgCl for the nucleosome arrays, and an oxygen scavenging system consisting of 0.01 units/ml of PCD, 2 mM of PCA, and 1 mM of aged Trolox to serve a Reducing Oxidizer System [50, 51].

### tFCS data recording and pre-processing

For each experiment, we collected data in continuous mode for 10-30 minutes, during which we recorded the fluorescence signal from the reference and probe dyes, the microscope stage trajectory, and the tracking laser power. These raw signals were semi-automatically pre-processed offline to detect and classify individual molecules, compute the fluorescence correlation functions, and correct for background effects (**SI Materials and Methods**)

### Compaction scores and clustering of conformational states

To compare molecular macrostates, tFCS data were quantified using a simple metric of molecular compaction. We defined the relative compaction between two molecules as the ratio of the inverse of the integral of the correlation function of the two molecules, computed over the interval of time-lags from 5 μs to 10 ms. These bounds were chosen empirically to encompass the region of the correlation signal which differs between the macrostates of interest, while avoiding introduction of noise from the shorter ( < 5 μs) and longer ( > 10 ms) timescales.

### DNA constructs labeling and nucleosome arrays reconstitution

DNA molecules used in all the experiments were fluorescently labeled with a single probe dye and a single reference dye by ligation or PCR. Nucleosome arrays were reconstituted using the salt dialysis method with purified histones CENP-A/H4 tetramers and H2A/H2B dimers as previously described [52], and using a 2.0 ratio of tetramer per 601 site (**SI Materials and Methods**).

## Contributions

H.M. and A.F.S. supervised the project. C.L., A.F.S. and H.M. designed the instrumentation and the method. C.L. built the instrument, did the tFCS experiments and processed the data. C.L., A.F.S. and H.M. designed and interpreted the DNA and chromatin dynamics experiments. C.L., J.C.B., A.F.S and H.M. designed and interpreted the LacI experiments. C.L. prepared the DNA and nucleosome arrays constructs, J.C.B. purified Lac repressor, and C.J.F. purified the recombinant histones. C.L. developed the analysis pipeline and wrote the code. C.L., J.C.B., A.F.S. and H.M. wrote and/or commented on the manuscript.

## Acknowledgments

We thank Michael Armen for his help with the instrumentation development and his technical assistance, and Andrew Spakowitz, Tom Lampo, Dmitri Pavlichin, Peter McMahon for insightful discussions on polymer dynamics and stochastic processes. We also thank Dodd Gray for his help with the manuscript, Whitney Johnson, Bradley French, Frederic Westhorpe for their feedback on the figures, and the members of the Mabuchi and Straight Labs for helpful discussions. This work was supported by NIH R01 GM106005 (to AFS and HM) and by NSF CMMI-0856205 (to HM).

